# Functional specialization of monocot DCL3 and DCL5 proteins through the evolution of the PAZ domain

**DOI:** 10.1101/2021.08.02.454693

**Authors:** Shirui Chen, Wei Liu, Masahiro Naganuma, Yukihide Tomari, Hiro-oki Iwakawa

**Author notes:** Correspondence (Y.T.), (H.-o.I.).

## Abstract

Monocot DICER-LIKE3 (DCL3) and DCL5 produce distinct 24-nt heterochromatic small interfering RNAs (hc-siRNAs) and phased secondary siRNAs (phasiRNAs). The former small RNAs are linked to plant heterochromatin, and the latter to reproductive processes. It is assumed that these DCLs evolved from an ancient “eudicot-type” DCL3 ancestor, which may have produced both types of siRNAs. However, how functional differentiation was achieved after gene duplication remains elusive. Here, we find that monocot DCL3 and DCL5 exhibit biochemically distinct preferences for 3′ overhangs and 5′ phosphates, consistent with the structural properties of their *in vivo* double-stranded RNA substrates. Importantly, these distinct substrate specificities are determined by the PAZ domains of DCL3 and DCL5 which have accumulated mutations during the course of evolution. These data explain the mechanism by which these DCLs cleave their cognate substrates from a fixed end, ensuring the production of functional siRNAs. Our study also indicates how plants have diversified and optimized RNA silencing mechanisms during evolution.

## Introduction

Small interfering RNAs (siRNAs) and microRNAs (miRNAs) are critical players in RNA silencing pathways which regulate various biological processes including organismal development and antiviral immunity^1–4^. These small RNAs are processed from either long double-stranded RNAs (dsRNAs) or RNAs with hairpin-like structures by specific ribonucleases called Dicer in animals or Dicer-like (DCL) proteins in plants^5,6^. These Dicer and DCL proteins are evolutionary conserved multidomain proteins belonging to the RNase III family^6^. While mammals have a single Dicer, plants encode multiple DCL proteins that produce different types of small RNAs^7^. For example, the genome of the model plant *Arabidopsis thaliana* encodes four DCL proteins, AtDCL1–4 with precise activities. AtDCL1 produces 20 to 22-nucleotide (nt) miRNAs, while AtDCL4 and 2 generate 21 and 22-nt siRNAs, respectively^7^. These small RNAs then regulate protein and mRNA levels through post-transcriptional gene silencing^7^. In contrast, AtDCL3 produces heterochromatic 24-nt siRNAs (hc-siRNAs) that form specific RNA-induced silencing complexes (RISCs) with ARGONAUTE4/6 (AGO4/6). RISCs promote sequence-specific DNA methylation and thus transcriptional gene silencing^8^. This RNA-directed DNA Methylation (RdDM) process is essential in repressing transposable elements, responding to stresses and maintaining genome integrity^9–11^. In short, the evolution of DCL proteins has led to diverse mechanisms that regulate gene expression at different levels.

AtDCL3 targets dsRNAs that are generated by the sequential action of two polymerases, DNA-dependent RNA polymerase IV (Pol IV) and RNA-dependent RNA polymerase 2 (RDR2)^12–15^. Pol IV synthesizes 30–40-nt RNAs (Pol IV strand), which often bear adenine at the 5′ end^16,17^. RDR2 then synthesizes the complementary strand of the Pol IV strand (RDR2 strand) through its RNA-dependent RNA polymerase activity^16^. The resulting dsRNAs are called Pol IV and RDR2-dependent RNAs (P4R2 RNAs)^18^. RDR2 tends to add one or two non-templated nucleotide(s) to the 3′ end of the RDR2 strand via its terminal nucleotidyl transferase activity. Thus, P4R2 RNAs typically harbor a 1- or 2-nt 3′ overhang on the RDR2 strand, while the 3′ end of Pol IV-generated strands are blunt-ended (Figure 1A**, left panel**)^16,18^. Interestingly, AtDCL3 preferentially cleaves P4R2 RNAs from the 5′ end of the Pol IV strand^17^. This asymmetric dicing by AtDCL3 is thought to be achieved by the combination of 5′ A selection upon Pol IV transcription and preference for the unstable 5′ A and U end resulting from AtDCL3 cleavage^17,19^. However, it remains unclear if the difference in thermodynamic stability alone explains the biased cleavage.

**Figure 1.**
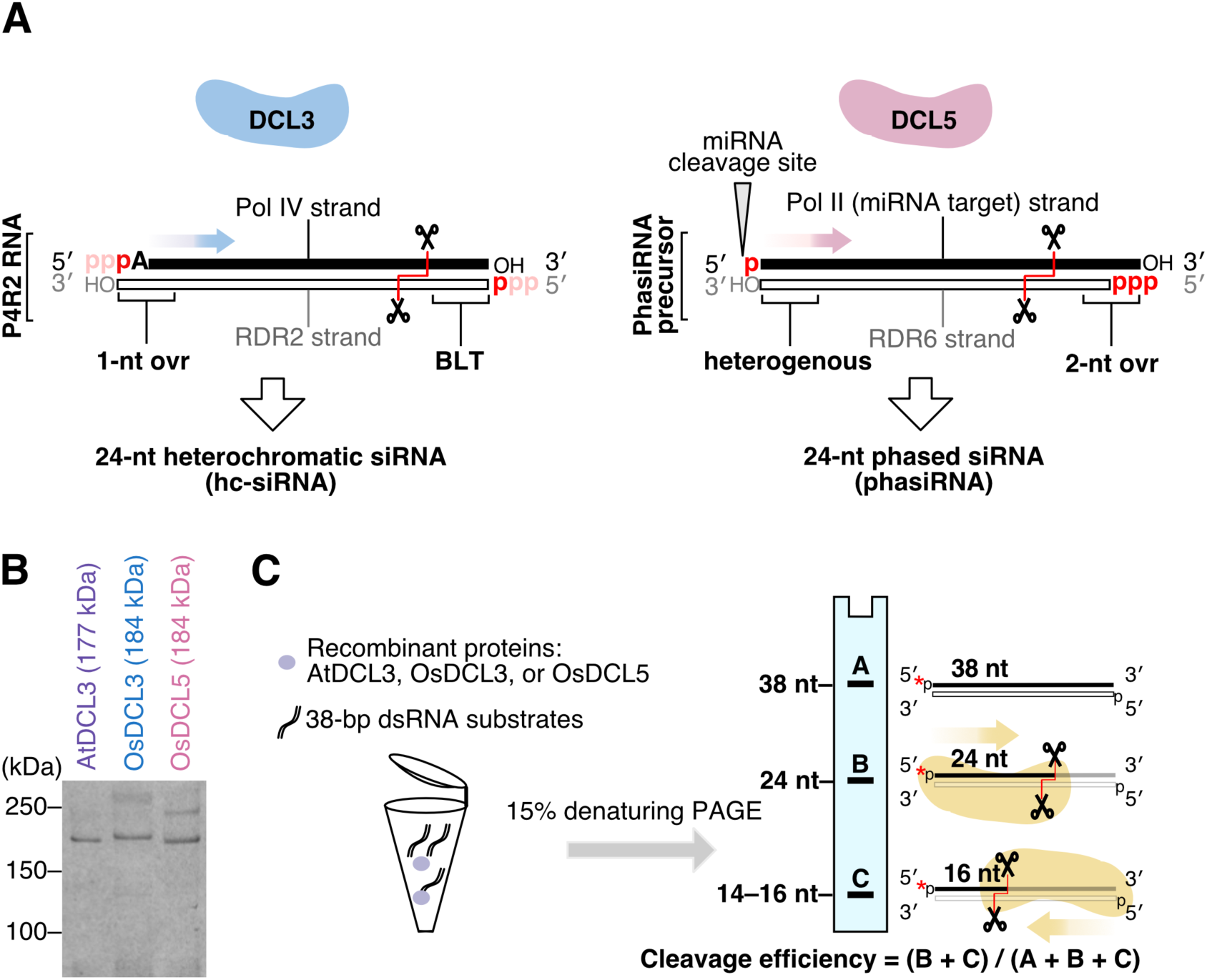
*In vitro* assay determines substrate specificities of DCL3 and DCL5. (A) Schematic illustrating *in vivo* substrates for DCL3 and DCL5. The dsRNA substrates for DCL3, named Pol IV and RDR2-dependent RNAs (P4R2 RNAs), have 5′ triphosphates on both strands in theory, which might be converted to monophosphates before cleavage by DCL3. The 3′ end structures of the two strands of P4R2 RNAs are different, and carry a 1-nt overhang (1-nt ovr) on one strand (RDR2 strand) and blunt end (BLT) on the other strand (Pol IV strand). The cleavage direction is shown with a blue arrow. DCL5 substrates are miR2275 targets, with dsRNA generated by RDR6. Such phasiRNA precursors generally carry heterogenous 3′ structures. The 5′ phosphorylation status of phasiRNA precursors is different on the two strands: with a 5′ monophosphate on one strand (miRNA target strand) and a 5′ triphosphate on the other strand (RDR6 strand). The cleavage direction is shown with a pink arrow. (B) Coomassie brilliant blue staining of recombinant AtDCL3, OsDCL3 and OsDCL5 proteins. (C) Schematic of *in vitro* dicing assays. dsRNA substrates can be cleaved from the 5′ ends of both strands, mainly generating two cleaved fragments: 24-nt and 14–16-nt. The products of dicing assays were analyzed by denaturing gel. Band A, B and C represent full-length RNA, 24-nt and 14–16-nt products, respectively. Cleavage ratio is calculated by dividing the sum of the cleaved fragments (B+C) by the total amounts of the substrate (A+B+C).

Although nascent transcripts of RNA polymerases generally have a triphosphate group at the 5′ end^15^, the P4R2 RNAs that accumulate in *dcl* mutant plants and mature hc-siRNAs both have a 5′ monophosphate group^17,18,20^. One explanation for this is that DCL3 cleaves P4R2 RNAs after the tri- to monophosphate conversion of the 5′ end through an unknown activity^17,21^. In contrast, a recent biochemical analysis showed that recombinant AtDCL3 cleaves not only substrates harboring a 5′ monophosphate, but also those with a 5′ triphosphate^16^. This suggests that AtDCL3 can directly cleave P4R2 RNAs with 5′ triphosphates *in vivo*.

Some plants produce 24-nt siRNAs that are distinct from hc-siRNAs^22^. These siRNAs are called reproductive phased secondary 24-nt siRNAs (24-nt phasiRNA), which are highly expressed in anthers^23,24^. Generally, phasiRNAs are produced from the RNAs targeted by 22-nt small RNAs^25–28^. The 22-nt small RNA-loaded Argonaute (AGO) proteins cleave the target RNA, resulting in the production of a 3′ cleavage fragment with a 5′ monophosphate. This fragment is then converted into a dsRNA with a triphosphate at the 5′ end of the antisense strand by RDR6, which is recruited via SILENCING DEFECTIVE 5 (SDE5) and the complex consisting of 22-nt small RNA, ARGONAUTE1 (AGO1) and SUPPRESSOR OF GENE SILENCING 3 (SGS3)^29^. Because RDR6 begins RNA synthesis at the third nucleotide of the template’s 3′ end^30^, the Pol II (sense) strand of the dsRNA has 2-nt 3′ overhang (Figure 1A**, right panel**). In contrast, the 3′ end of the RDR6 (antisense) strand of the dsRNA is more heterogeneous, having a blunt end or bearing a 1-nt or 2-nt non-templated nucleotide added by the terminal nucleotidyl transferase activity of RDR6 (Figure 1A**, right panel**)^30–32^. The dsRNA intermediate is then processed by DCLs into phasiRNAs, with the phase determined by the small RNA-guided cleavage site. The mechanism of this one-way processing remains unclear. In monocots, DCL5, which is thought to have evolved via duplication of DCL3 (also called DCL3b), specifically produces 24-nt phasiRNAs in the anther (Supplementary Figure 1)^33^. Although eudicots are believed to lack the 24-nt phasiRNA pathway, recent studies argue that some eudicots like *Citrus sinensis* and *Populus trichocarpa* produce 24-nt phasiRNAs even without encoding DCL5 (Supplementary Figure 1)^22,28^. In these plants, DCL3 needs to produce both 24-nt phasiRNAs and hc-siRNAs. Given the completely different structures of the dsRNA precursors of 24-nt phasiRNAs and hc-siRNAs, DCL5 and monocot and eudicot DCL3 need to tailor their substrate specificities to their cognate targets. However, so far, the biochemical properties of these DCLs, which are relevant to gene regulation and reproduction, have not been examined and compared.

Dicer and DCL proteins generally consist of five functional domains: the helicase domain, PAZ (PIWI, AGO, and Zwille) domain, two RNase III domains and double-stranded RNA-binding domain from N to C terminal^6,34,35^. Previous biochemical and structural studies of human DICER1 and *Drosophila melanogaster* Dicer-2 (Dcr-2) demonstrated that the PAZ domain has two pockets that bind the 5′ and 3′ ends of the substrate dsRNA respectively. These binding pockets are critical for the precise production of small RNAs^36–38^. In addition to the PAZ domain, it is reported that the helicase domain interacts with the substrate dsRNA and is required for *Drosophila* Dcr-2 to bind the 3′ end^39–41^. However, because the sequences of PAZ and helicase domains of plant DCLs greatly differ from human Dicer and *Drosophila* Dcr-2^42^, the mechanisms underlying substrate recognition and specific catalysis by plant DCLs remain unclear.

In this study, we succeeded in preparing fully functional recombinant eudicot AtDCL3, monocot *Oryza sativa* DCL3 (OsDCL3) and DCL5 (OsDCL5). Our analysis elucidates how DCL3 and DCL5 have become functionally specialized after gene duplication. OsDCL5 and OsDCL3 have distinct substrate specificities for both 5′ phosphate and 3′ structures, reflecting the different *in vivo* functions of these proteins. Moreover, we find that the PAZ domain is a key determinant of DCL3 and DCL5 substrate specificity. These preferences explain how DCL3 and DCL5 cleave substrates from the fixed end to ensure the production of functional siRNAs. Importantly, eudicot AtDCL3 has intermediate preferences between OsDCL5 and OsDCL3 for both a blunt ended structure and 5′ triphosphate. This argues that OsDCL5 and OsDCL3 became functionally specialized after duplication from the “eudicot-type” DCL3. Taken together, our study provides insights into the functional differentiation of DCLs via the evolution of the PAZ domains. This provides a molecular understanding of how plants have diversified and optimized RNA silencing mechanisms through DCL gene duplication.

## Results

### DCL5 and DCL3 proteins have different preferences for 3′ dsRNA structures

To compare the substrate preferences of monocot DCL5 and monocot and eudicot DCL3 proteins *in vitro*, we successfully prepared full-length recombinant DCL proteins: OsDCL5, OsDCL3, and AtDCL3 using *Drosophila* S2 cells (Figure 1B). Double-stranded RNA (dsRNA) substrates were radiolabeled at the 5′ end of the sense or antisense strand and incubated with purified recombinant DCL for the *in vitro* dicing assay. Each strand of the substrate was 38 nt long, thus mimicking the length of P4R2 RNAs. Since dsRNAs can be cleaved from both ends, two product bands (24-nt and ∼16-nt) were expected (Figure 1C). The cleavage efficiency was calculated by dividing the sum of the cleaved fragments by the total amount of the substrate (full-length + cleaved fragments). To determine the 3′ structures preferred by DCL3 and DCL5 proteins respectively, we performed *in vitro* dicing assays with dsRNAs harboring different 3′ structures: blunt end (BLT), 1-nt overhang (1ovr), and 2-nt overhang (2ovr) (Figure 2A). Both AtDCL3 and OsDCL3 cleaved dsRNAs with overhangs more efficiently than the BLT substrates (Figure 2B, C). At early time points (5–30 minutes after incubation), OsDCL3 showed a more pronounced preference for overhangs than AtDCL3 (Figure 2B, C). In contrast, OsDCL5 cleaved BLT, 1ovr and 2ovr dsRNA substrates with similar efficiency (Figure 2D), showing no specific preference for 3′ structures. Interestingly, both AtDCL3 and OsDCL3 generated multiple cleavage products from BLT substrates (Figure 2B, C). We speculate that AtDCL3 and OsDCL3 cannot accurately process BLT substrates, resulting in intermediate products that are longer than 24-nt. These intermediate products might then be cleaved again, generating fragments shorter than 16-nt (Figure 2B, C). In contrast, OsDCL5 cleaved BLT as accurately as 1ovr and 2ovr substrates. Taken together, we conclude that DCL5, monocot and eudicot DCL3 proteins have different preferences for the 3′ structure of dsRNAs; OsDCL3 has the strongest preference for 3′ overhangs, followed by AtDCL3, while OsDCL5 has no apparent preference for specific 3′ structures.

**Figure 2.**
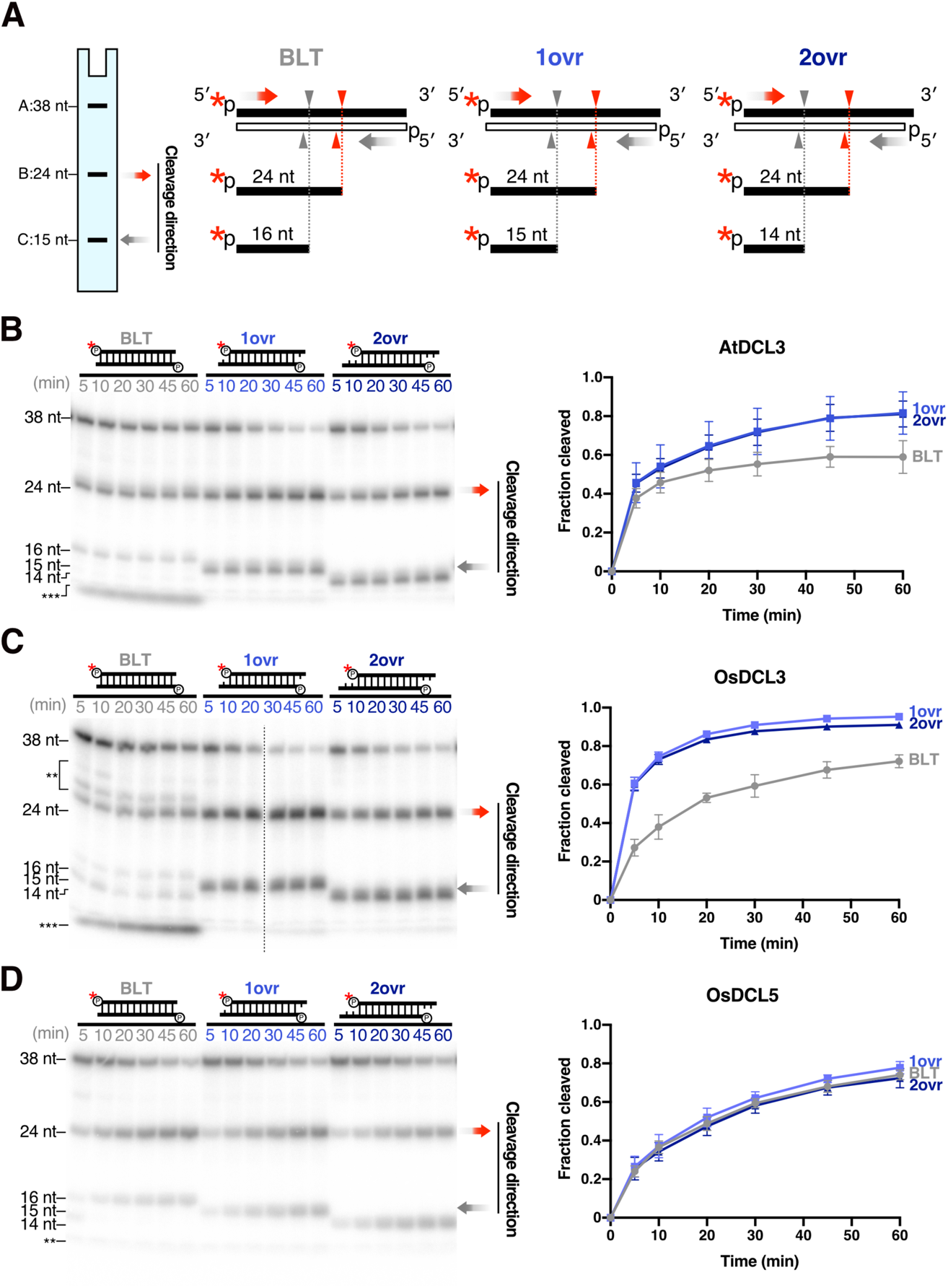
DCL3 and DCL5 have different preferences for 3′ structures. (A) 38-nt substrates radiolabeled on the 5′ monophosphate of sense strands, with different 3′ structures: blunt (BLT), 3′ 1-nt overhang (1 ovr) and 3′ 2-nt overhang (2 ovr), were used in the dicing assays. Red arrowheads and grey arrowheads indicate cleavage from the 5′ end of sense strands and antisense strands, respectively. These substrates can be cleaved into 24-nt (band B) and 14,15 or 16-nt (band C) products according to their 3′ structures. Cleavage efficiency is calculated using (B+C)/(A+B+C). (B, C, D) Left panel: Representative gel images of dicing assays by (B) AtDCL3, (C) OsDCL3 and (D) OsDCL5 cleaving the 38-nt substrates with different 3′ structures (from left to right: BLT, 1ovr and 2ovr). The double and triple asterisks indicate the inaccurate cleavage products, long and short, respectively. Right panel: Quantification of cleavage efficiency in the left panel. For BLT, bands of inaccurate cleavage products (long and short fragments) were also calculated as cleavage products. The mean values ± SD from three independent experiments are shown. AtDCL3 and OsDCL3 both prefer substrates with 3′ overhangs. OsDCL5 does not show preferences for specific 3′ structures.

### The 5′ phosphate of dsRNAs is required for efficient cleavage by both DCL3 and DCL5

In addition to the recognition of the 3′ structure, the recognition of the 5′ end of RNA substrates is also important for both accurate and efficient dicing of dsRNAs^39,43^. Previous *in vitro* dicing assays using crude plant lysates confirmed that a 5′ phosphate is required for AtDCL3-mediated cleavage of dsRNAs carrying 3′ overhangs^19^. To investigate the importance of the 5′ phosphate of dsRNAs in DCL3- and DCL5-mediated cleavage, we performed *in vitro* dicing assays with 3′ 1-nt overhang substrates radiolabeled at the 5′ monophosphate of antisense strands. These substrates carry either a 5′ monophosphate group (MonoP) or a hydroxyl group (OH) on the sense strand (Figure 3A). If the 5′ monophosphate is required for substrate processing, a 5′-hydroxyl should decrease the generation of 15-nt cleavage products which arise from the 5′ end of the sense strand. We found that, for all three DCL proteins, 15-nt products generated from a 5’-OH substrate were decreased compared to MonoP substrates (Figure 3B–D. This result argues that the 5′ phosphate of the substrate is required for efficient dsRNA cleavage by both DCL3 and DCL5.

**Figure 3.**
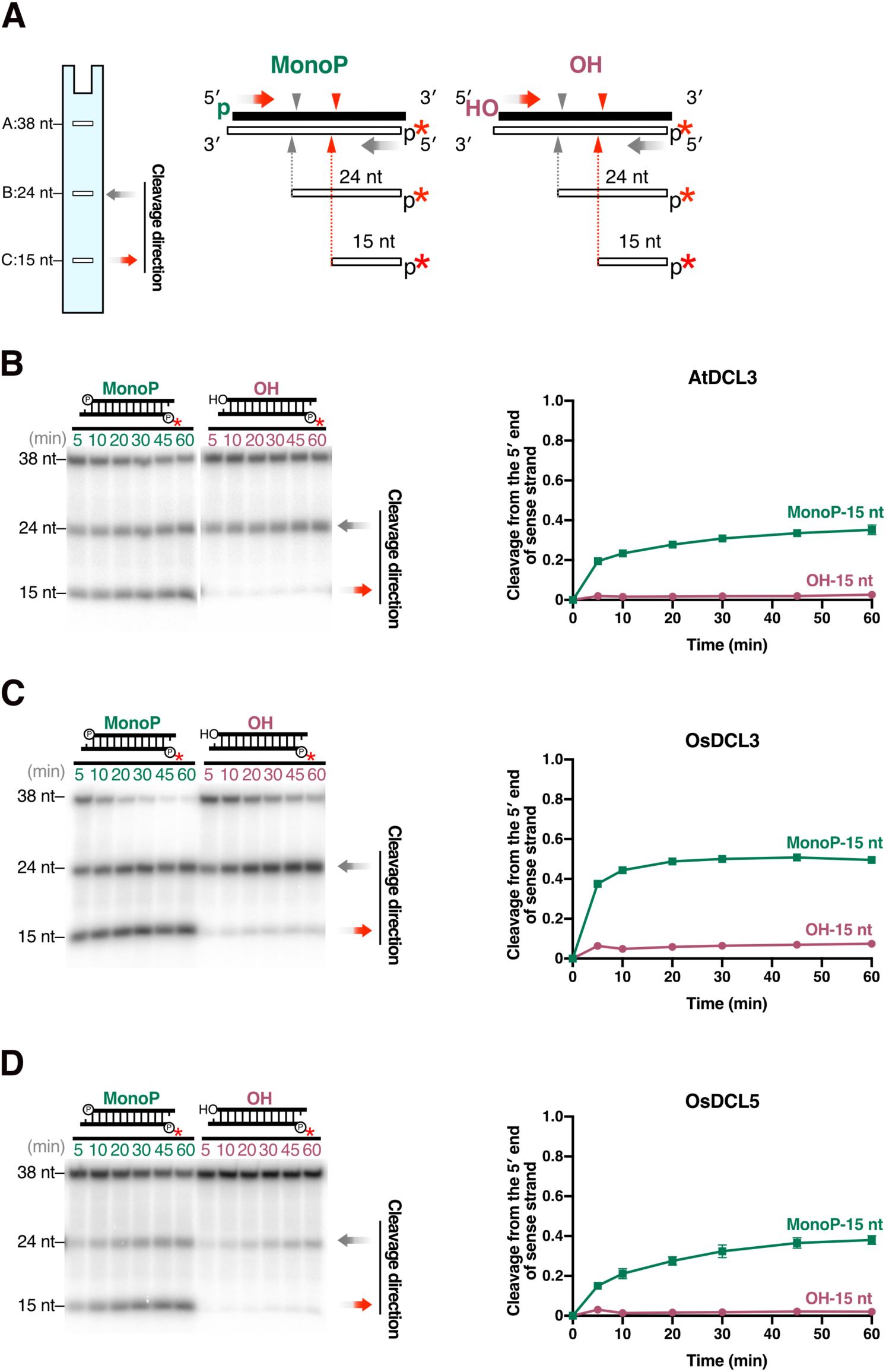
The 5′ monophosphate is important for catalysis by DCL3 and DCL5. (A) 1-nt 3′ overhang substrates radiolabeled on the 5′ monophosphate of antisense strands, with a 5′ hydroxyl groups (OH) or a monophosphate (MonoP) on sense strands, were used for dicing assays. Red arrowheads and grey arrowheads indicate cleavage from the 5′ end of sense strands and antisense strands, respectively. Cleavage from the 5′ end of sense strands results in 15-nt products (band C), and the proportion of cleavage from the 5′ end of sense strands is calculated by C/(A+B+C). (B, C, D) Left panel: Representative gel images of dicing assays by (B) AtDCL3, (C) OsDCL3 and (D) OsDCL5 cleaving 1-nt overhang MonoP and OH substrates. Right panel: Quantification of cleavage from the 5′ end of sense strands (15-nt bands) in the left panel. The mean values ± SD from three independent experiments are shown. Compared with a 5′ monophosphate, a 5′ hydroxyl group greatly reduced cleavage from the 5′ end of sense strand by AtDCL3, OsDCL3 and OsDCL5.

### DCL3 and DCL5 have distinct preferences for a 5′ triphosphate on the dsRNA

In theory, newly synthesized RNAs generated by Pol IV and RDR2 carry 5′ triphosphates. It is therefore possible that P4R2 RNAs carry a 5′ triphosphate when they encounter DCL3. In addition, precursors of phasiRNAs, i.e. DCL5 substrates, are also likely to possess a triphosphate group at the 5′ end of the antisense strand, which is synthesized by RDR6. To investigate the effect of a 5′ triphosphate group DCL3 and DCL5-mediated cleavage, we performed *in vitro* dicing assays with dsRNA substrates carrying a 5′-^32^P on the antisense strand. Substrates were monophosphorylated (MonoP) or triphosphorylated (TriP) at the 5′ end of their sense strands. Since cleavage from the 5′ end of the sense strands results in 15-nt products, the preference for the 5′ phosphate can be quantitated by comparing the proportion of 15-nt bands generated from TriP and MonoP substrates (Figure 4). We found that OsDCL5 generated a lower proportion of 15-nt product from TriP compared to MonoP substrates (Figure 4D), indicating that the 5′ triphosphate group strongly inhibits OsDCL5-mediated cleavage. Similarly, the 5′ triphosphate also negatively affected dicing by AtDCL3, although the effect was less obvious than that on OsDCL5 (Figure 4B). In contrast, the proportion of 15-nt products cleaved by OsDCL3 was similar for MonoP and TriP substrates (Figure 4C). Thus, the 5′ triphosphate does not affect OsDCL3-mediated dsRNA cleavage. In conclusion, DCL5 and DCL3 proteins have different cleavage efficiencies based on the triphosphate group at the 5′ end of dsRNA, likely impacting the small RNA substrates and pathways they can act upon in plants.

**Figure 4.**
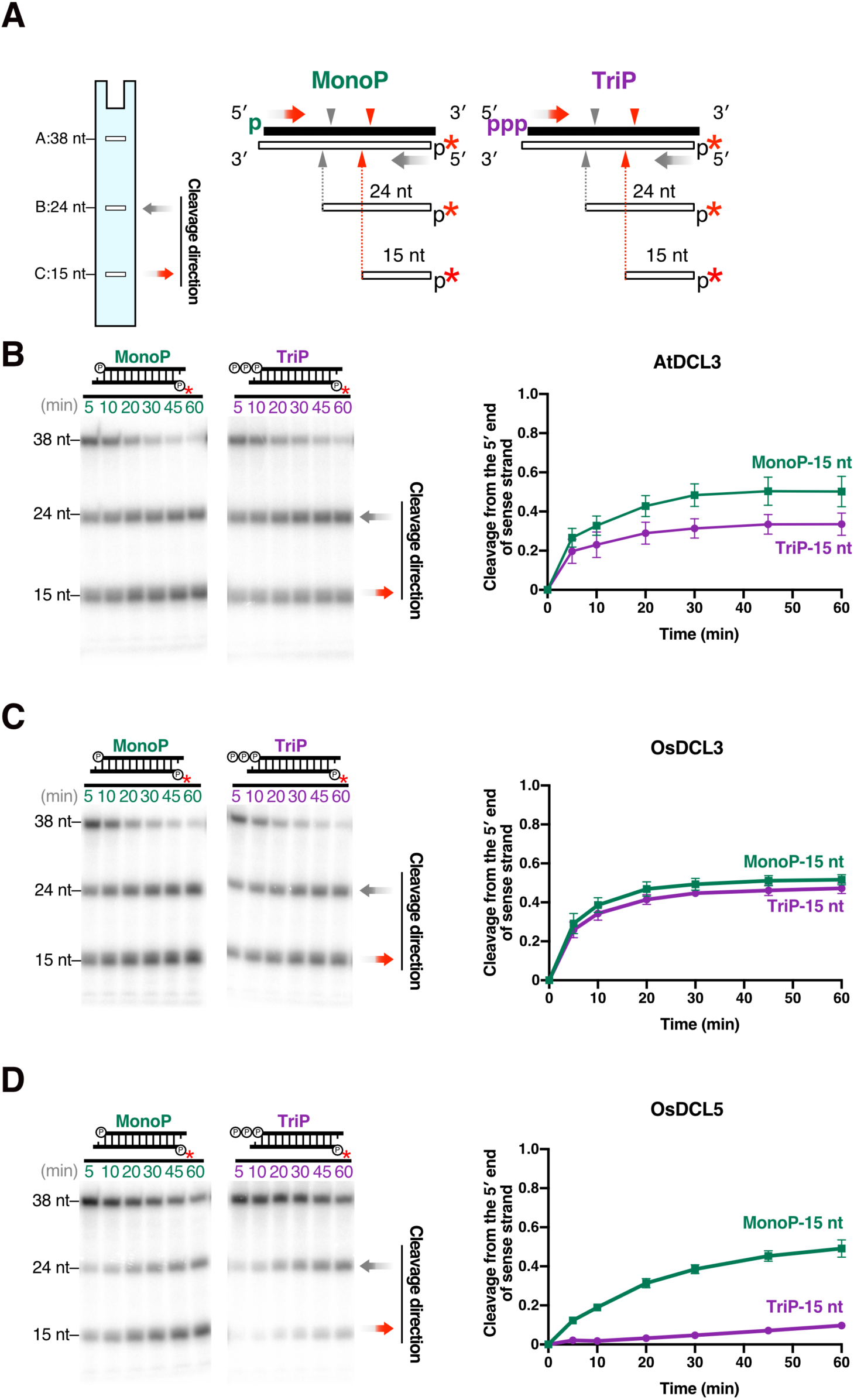
DCL3 and DCL5 have different preferences for the 5′ triphosphate. (A) Dicing assays were conducted using 1-nt 3′ overhang substrates radiolabeled on the 5′ monophosphate on the antisense strand, carrying a 5′ monophosphate (MonoP) or triphosphate (TriP) on the sense strands. Red arrowheads and grey arrowheads indicate cleavage from the 5′ end of sense strands and antisense strands, respectively. Cleavage from the 5′ end of sense strands results in 15-nt products (band C), and the proportion of cleavage from the 5′ end of sense strands is calculated as C/(A+B+C). (B, C, D) Left panel: Representative gel image of dicing assays with (B) AtDCL3, (C) OsDCL3 and (D) OsDCL5 cleaving MonoP and TriP substrates. Right panel: Quantification of the proportion of cleavage from the 5′ end of sense strands (15-nt bands) in the left panel. The mean values ± SD from three independent experiments are shown. AtDCL3 slightly prefers substrates with 5′ monophosphate. OsDCL3 does not show an obvious preference for the 5′ mono- or triphosphate. OsDCL5 prefers substrates carrying a 5′ monophosphate.

### The PAZ domain determines DCL cleavage preferences based on the dsRNA 3′ structure

Previous studies showed that the PAZ domain of Dicer proteins determines recognition of the 3′ dsRNA structure, in humans and *Drosophila*^37,38,44,45^. The strong preference of DCL3 for the 3′ overhang prompted us to hypothesize that the interaction between the PAZ domain and the 3′ end of dsRNAs is required for DCL3-mediated dsRNA cleavage. To test this, we introduced an extra phosphate group at the 3′ end of the antisense strand of the 1ovr substrate (1ovr 3′ p) (Figure 5A). This modification is expected to sterically block accommodation of the 3′ overhang by the PAZ domain (Supplementary Figure 2). If the 3′ phosphate inhibits substrate binding to the PAZ domain of DCLs, 24-nt fragments, i.e. cleavage products from the 5′ end of the sense strand, should decrease. In contrast, 15-nt fragments, which represent cleavage from the 5′ end of antisense strand, should increase. Our *in vitro* dicing assays with AtDCL3 or OsDCL3 showed a drastic decrease in the 24-nt fragment and increase in the 15-nt fragment when the 1ovr 3′ p substrate is cleaved. This indicates that 3′ end recognition is important for dicing by AtDCL3 and OsDCL3 (Figure 5B, C). In contrast, the proportion of 24-nt and 15-nt products generated by OsDCL5 was not greatly altrered by the addition of a 3′ phosphate to the antisense strand (Figure 5D). These data argue that the 3′ end of dsRNA is not strictly recognized by the PAZ domain of DCL5.

**Figure 5.**
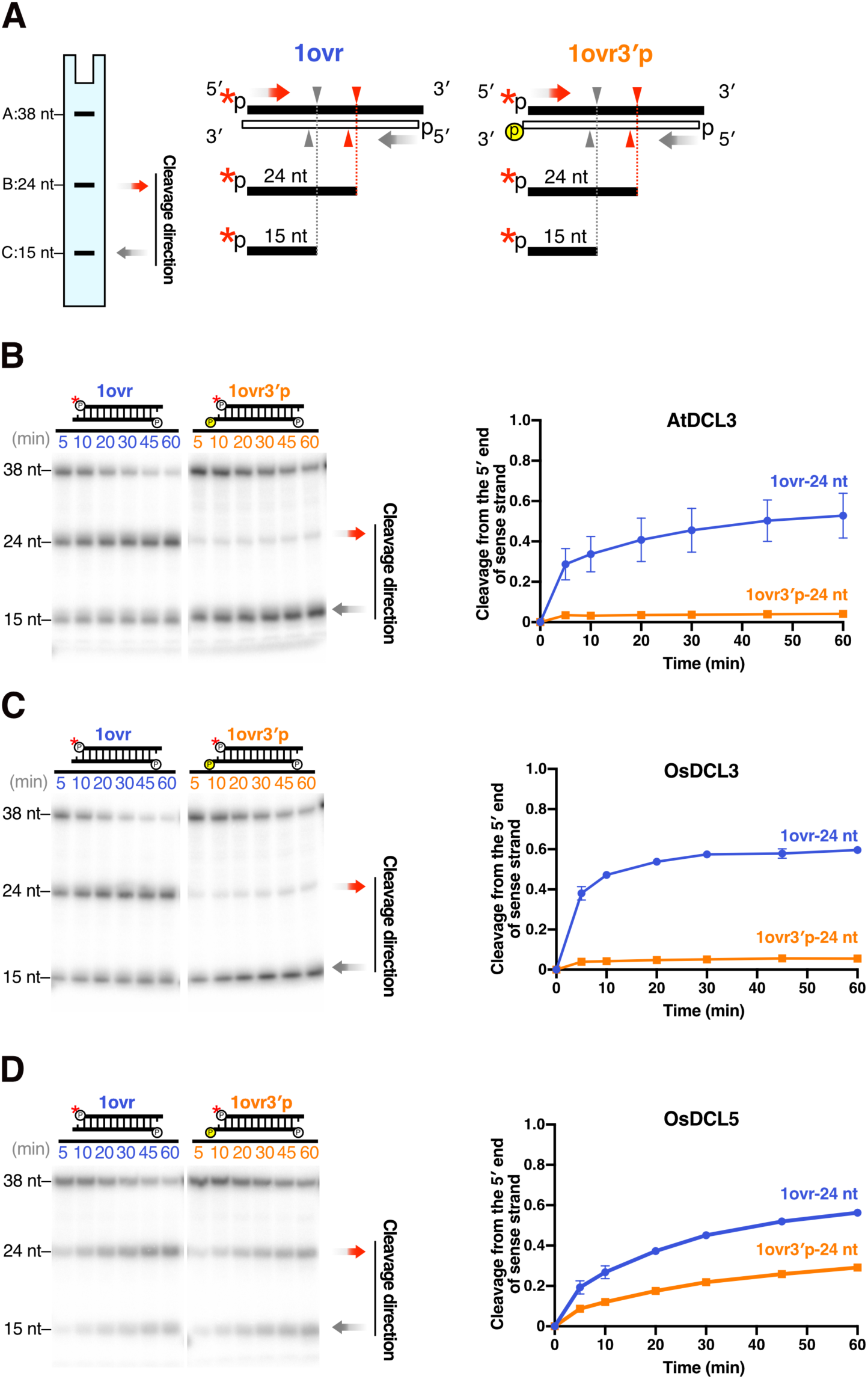
Recognition of the 3′ hydroxyl group is important for DCL3, but not DCL5, cleavage. (A) Dicing assays were conducted on 1-nt 3′ overhang substrates radiolabeled on the 5′ monophosphate of sense strands (1ovr). To disrupt recognition of the 3′ hydroxyl, an extra monophosphate group was added to the 3′ end of the antisense strand (1ovr 3′p). Red arrowheads and grey arrowheads indicate cleavage from the 5′ end of sense strands and antisense strands, respectively. Cleavage from 5′ end of sense strands (corresponding to the 3′ phosphate on antisense strands) results in 24-nt products (band B). (B, C, D) Left panel: Representative gel images of dicing assays by (B) AtDCL3, (C) OsDCL3 and (D) OsDCL5 cleaving 1-nt overhang substrates with or without a 3′ monophosphate on the antisense strand (1ovr 3′p or 1ovr). Right panel: Quantification of the proportion of cleavage from the 5′ end of sense strands (24-nt bands) is shown in the left panel. Mean values ± SD from three independent experiments are shown. The extra 3′ phosphate greatly reduced cleavage from the 5′ end of sense strand by AtDCL3 and OsDCL3 compared with the 1ovr substrate, but only moderately reduced cleavage from the 5′ end of sense strand by OsDCL5.

To further confirm the importance of the PAZ domain for 3′ recognition, we swapped the PAZ domains between OsDCL3 and OsDCL5 (Figure 6A). We named these chimeric proteins OsDCL3_PAZ5 and OsDCL5_PAZ3, and performed *in vitro* dicing assays to investigate their preferences for 3′ structures and 5′ phosphate. As with OsDCL5, we found that OsDCL3_PAZ5 prefers BLT substrates as well as substrates with 3′ overhangs (Figure 2D and 6B). In contrast, OsDCL5_PAZ3 showed a slight preference for substrates with 3′ overhangs, like OsDCL3 (Figure 2C and 6C). In addition, we observed that OsDCL3_PAZ5 cleaved BLT substrates as accurately as OsDCL5 (Figure 2D and 6B), whereas OsDCL5_PAZ3 produced multiple bands, like OsDCL3 (Figure 2C and 6C). We also performed dicing assays using substrates with or without an extra 3′ phosphate on the antisense strand (1ovr 3′p vs. 1ovr). Swapping PAZ domains between OsDCL3 and OsDCL5 reversed the effects of blocking 3′ end recognition on each protein (Supplementary Figure 3). Taken together, we conclude that the PAZ domain plays an important role in determining the preference for 3′ structure and cleavage fidelity of substrates with blunt ends.

**Figure 6.**
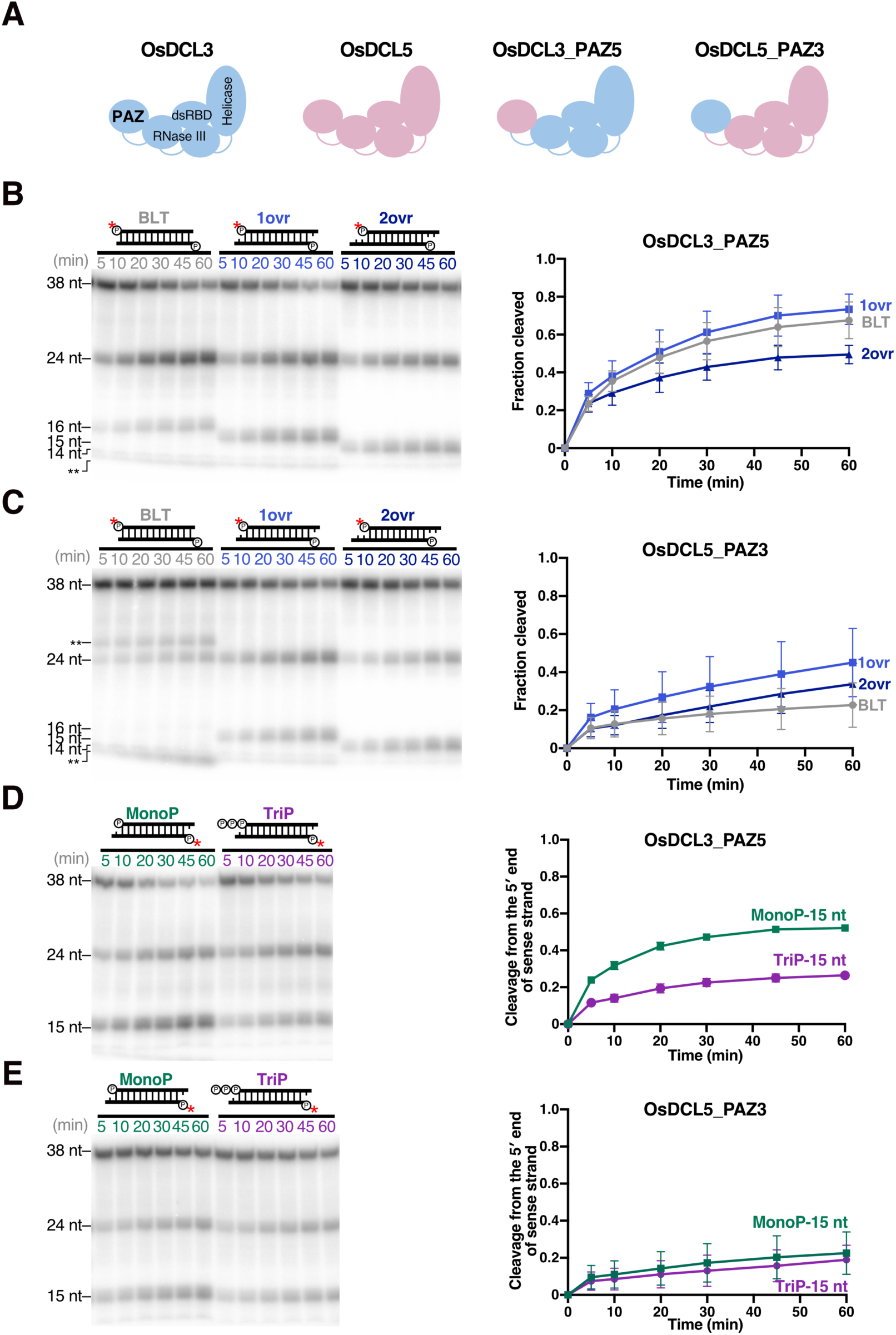
The PAZ domains plays an important role in specific DCL3 and DCL5 substrate preferences. (A) Swapping the OsDCL3 and OsDCL5 PAZ domains generates two chimeric proteins: OsDCL3_PAZ5 and OsDCL5_PAZ3. (B, C) Dicing assays by (B) OsDCL3_PAZ5 and (C) OsDCL5_PAZ3 with dsRNA substrates holding different 3′ structures. Left panel: Representative gel images of dicing assays by OsDCL3_PAZ5 and OsDCL5_PAZ3 cleaving 38-nt substrates BLT, 1ovr and 2ovr. The double and triple asterisks indicate the inaccurate cleavage products, long and short, respectively. Right panel: Quantification of dicing efficiency in the left panel. For BLT, bands of inaccurate cleavage products (long and short fragments) were also calculated as cleavage products. Mean values ± SD from three independent experiments are shown. OsDCL3_PAZ5 does not show an obvious preference for specific 3′ structures, while OsDCL5_PAZ3 prefers substrates with a 3′ overhang rather than blunt ends. (D, E) Dicing assays by (D) OsDCL3_PAZ5 and (E) OsDCL5_PAZ3 with MonoP and TriP dsRNA substrates. Left panel: Representative gel images of dicing assays by OsDCL3_PAZ5 and OsDCL5_PAZ3 cleaving 38-nt substrates, MonoP and TriP. Right panel: Quantification of the proportion of cleavage from the 5′ end of sense strands (15-nt bands) in the left panel. The mean values ± SD from three independent experiments are shown. OsDCL3_PAZ5 prefers 5′ monophosphate, while OsDCL5_PAZ3 does not discriminate 5′ monophosphate from triphosphate.

### The PAZ domain in DCL5 and DCL3 proteins determines 5′ phosphate preference on dsRNA substrates

Previous studies have demonstrated that, in human Dicer, several basic amino acid residues in the “core” of the PAZ domain and its upstream Platform domain form a binding pocket for the 5′ end of the dsRNA substrate^38,43^. Although the exact positions of these residues are not conserved in *Drosophila* Dcr-2, alternative basic residues within and upstream of the core PAZ domain are thought to form the binding pocket for the 5′ phosphorylated end of dsRNAs^36^. Since DCL3 and DCL5 lack corresponding basic residues upstream of the core PAZ domain (Supplementary Figure 4A), it remains unclear which domain(s) discriminate the 5′ monophosphate from the 5′ triphosphate of dsRNAs. Strikingly, we discovered that swapping the PAZ domains between OsDCL3 and OsDCL5 precisely altered 5′ phosphate preference. OsDCL3_PAZ5 produced less 15-nt fragments from TriP than from MonoP (Figure 6D), indicating that OsDCL3_PAZ5 mimics OsDCL5 and prefers a 5′ monophosphate. In contrast, OsDCL5_PAZ3 produced similar amounts of 15-nt fragments from the MonoP and TriP substrates (Figure 6E), indicating that, like OsDCL3, OsDCL5_PAZ3 does not discriminate 5′ monophosphate from triphosphate. Taken together, swapping the PAZ domain alters the preference for 5′ triphosphate in DCL5 and DCL3 proteins. These results suggest that, in addition to the 3′ structure, the PAZ domains of DCL3 and DCL5 determine 5′ phosphate preference during dsRNA cleavage. The PAZ domain therefore plays a key role in determining substrate preference in plant small RNA-mediated silencing pathways.

## Discussion

### Role of PAZ domains in determining DCL3 family substrate preferences

Previous studies have proposed that substrate preferences of human Dicer and *Drosophila* Dcr-2 proteins are determined by the PAZ domain and helicase domain^36,38,39,43,46^. In our study, swapping the PAZ domain was sufficient to alter substrate preferences for both 3′ structures and 5′ triphosphates for OsDCL3 and OsDCL5. Our data demonstrate that the PAZ domain alone can determine which dsRNA ends are preferred in DCL3 family proteins. By comparing the amino acid sequences of PAZ domains across DCL3 family proteins, we identified a variable region where monocot DCL3 and DCL differ (Supplementary Figure 4B). The corresponding region of human Dicer is located between the 5′ and the 3′ binding pockets in the platform-PAZ cassette. The cassette forms two structurally distinct complexes with short dsRNAs^38^; one has a visible *α*-helix that separates the two pockets, with the 3′ end of the dsRNA anchored in the 3′ pocket and the 5′ end released from the 5′ pocket (Supplementary Figure 4C); the other has a disrupted *α*-helix that allows anchoring of both ends of the dsRNA in the two pockets. Thus, the *α*-helix is directly linked to substrate dsRNA binding. Given that the amino acid sequences corresponding to the *α*-helix in human Dicer differ significantly between monocot DCL3 and DCL5 (Supplementary Figure 4B), these sequences are expected to form distinct 5′ and 3′ binding pockets, which may impact dsRNA recognition in monocot DCL3 and DCL5. Future structural analysis should reveal the exact mechanism by which the PAZ domains of DCL3 and DCL5 recognize different substrates.

### An additional rule for the biased processing of 24-nt heterochromatic siRNAs by DCL3

Pol IV collaborates with RDR2 to generate ∼37-nt long dsRNA precursors, P4R2 RNAs, for hc-siRNA production^17^. Interestingly, most of the sequenced mature hc-siRNAs are produced from the 5′ end of the Pol IV strand^17^. Currently, this bias is thought to be established by the preference for a 5′ adenine at the three different steps in the hc-siRNA biogenesis: (1) start site selection by Pol IV^17^ , (2) determination of the dicing direction by DCL3^19^, and (3) guide strand selection by AGO4/6^47^. In addition to the 5′A preference, we found that the recognition of the 3′ end structure of the dsRNA precursor by DCL3 is important for the establishment of substrate bias. A recent study showed that the precursor dsRNA for hc-siRNAs has an asymmetric structure, with a 1-nt overhang at one end and a blunt structure at the other end^16^. Our study with recombinant proteins and a previous study with a crude cytoplasmic extract demonstrated that DCL3 prefers 3′ overhangs to blunt-ended structures (Figure 2B, 2C)^19^. This substrate specificity will allow DCL3 to preferentially cleave the dsRNA precursor from the 5′ end of the Pol IV strand. This strand forms a 1-nt 3′ overhang structure with the 3′ end of the RDR2 strand (Supplementary Figure 5A**, left**)^16^. Cleavage from the opposite end would be less efficient because of its blunt-ended nature (Supplementary Figure 5A**, left**)^16^. This 3′ structure bias would further promote the biased production of 5′ adenine 24-nt siRNAs together with the 5′ A preferences by Pol IV and DCL3. These multi-layer selection steps may facilitate the formation of AGO4/6-RISC, resulting in efficient RNA-directed DNA methylation.

### Preference for a 5′ monophosphate in OsDCL5 determines the direction of cleavage for phased 24-nt siRNA production

Specific miRNAs, including miR390 and 22-nt small RNAs, recruit RDR6 to the target RNA to generate dsRNA precursors^29^. This long dsRNA is then processed into phasiRNAs by DCLs^28^. Interestingly, DCLs always cleave precursors from the miRNA-mediated cleavage site toward the other end. Although this fixed orientation of dicing is important for production of functional phasiRNAs, how this is achieved has remained unclear. In this study, we found that DCL5, which is known to produce phasiRNAs from RNAs with 22-nt miR2275 target sites, cleaves dsRNAs from a 5′ monophosphate much more efficiently than a 5′ triphosphate. Since the miRNA-cleaved end has a 5′ monophosphate, whereas the 5′ end of the RDR6 strand has a triphosphate in theory, DCL5 substrate preference explains the directionality of dicing (Supplementary Figure 5A**, right**).

Although *Arabidopsis* does not produce 24-nt phasiRNAs, some eudicots do so even without encoding DCL5^22,28^. In these plants, DCL3 is likely to be responsible for generating 24-nt phasiRNAs. We envision that the slight preference of eudicot DCL3 for 5′ monophosphorylated ends may also contribute to directional processing to produce functional phasiRNA precursors.

### The preference for 5′ triphosphates in OsDCL3 and AtDCL3 may enhance the production of heterochromatic siRNAs

Since Pol IV uses nucleoside triphosphates (NTPs) as substrates for transcription and lacks binding regions for the capping complexes, Pol IV-synthesized transcripts are expected to possess a triphosphate group at the 5′ end^15,21^. RDR2 also generates 5′ triphosphate RNAs *in vitro*^15,16^. Thus, nascent P4R2 RNAs theoretically possess 5′ triphosphates at both ends. However, previous studies showed that the P4R2 RNAs that accumulate in the *dcl2/3/4* mutant have monophosphates at the 5′ ends^17,18,20^, raising the possibility that unknown RNA phosphatases convert the 5′ triphosphates of dsRNAs into monophosphates in nuclei. In wild-type plants, DCL3 may encounter the P4R2 RNAs before or after the tri- to monophosphate conversion. In any case, OsDCL3’s ability to cleave both 5′ mono- and triphosphorylated dsRNAs with the same efficiency will maximize the production of 24-nt heterochromatic siRNAs (Figure 5C **and** Supplementary 5B). Given that AtDCL3 has a slight preference for 5′ monophosphorylated over triphosphorylated precursors (Figure 5B), dephosphorylation prior to dicing may enhance the production of heterochromatic 24-nt siRNAs in *Arabidopsis thaliana*.

### Functional specialization of duplicated DCL3 genes in monocots

It is now believed that the appearance of DCL5 in monocots is explained by the “sub-functionalization” of the ancestral DCL3 gene, which is speculated to function in the production of both hc-siRNAs and phasiRNAs^28^. However, there is no biochemical evidence supporting this hypothesis. One of the most interesting results in our study may be that monocot OsDCL3 and OsDCL5 have completely different substrate specificities, whereas eudicot AtDCL3 has an intermediate preference for dsRNAs with a 5′ triphosphate and 3′ overhang structure. This implies that monocot DCL5 and DCL3 were not only subfunctionalized, but further optimized for cognate substrates after the duplication from the ancient “eudicot-type” DCL3 (Figure 7). This functional specialization process appears to have been achieved through accumulation of mutations in the PAZ domain. Further biochemical studies on DCLs in a wider variety of plant species will reinforce this hypothesis. Our data, however, indicate how OsDCL family members have evolved to function in specific biological pathways.

**Figure 7.**
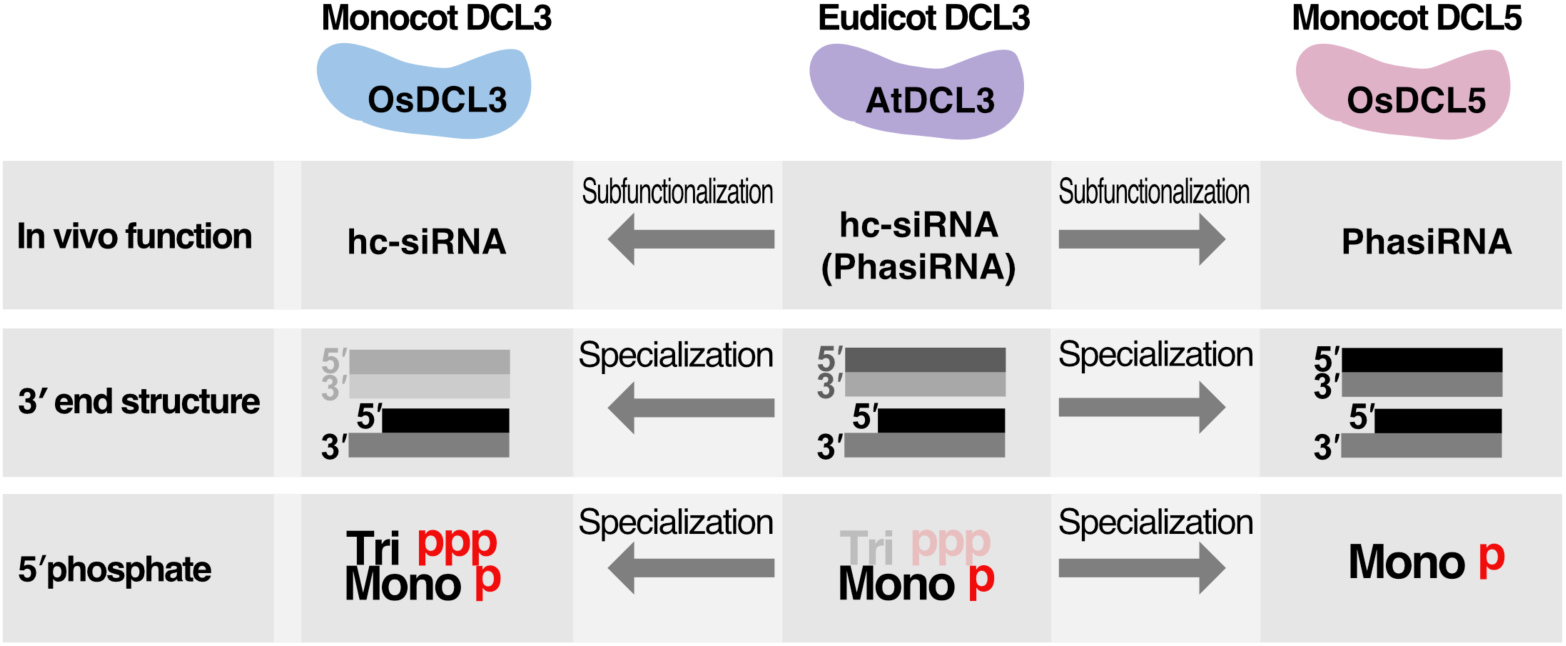
Summary of functions and substrate preferences of DCL3/5 family proteins. Summary of substrate functions and specificities of eudicot DCL3, monocot DCL3 and DCL5. Monocot OsDCL3 and OsDLC5 have completely distinct substrate specificities, whereas eudicot AtDCL3 has an intermediate preference for dsRNAs. These data argue that monocot DCL5 and DCL3 duplicated from a common ancestral “eudicot-type” DCL3, then subfunctionalized and further optimized to cleave cognate substrates.

## Materials and Methods

### Plasmid construction

The primers used in this study are listed in Supplementary Table S1.

#### pASW-AtDCL3

AtDCL3 ORF was amplified from *Arabidopsis thaliana* (Col-0) cDNA using Oligo No. 1 and 2, and cloned into TOPO^®^ vector using pENTR™/D-TOPO™ Cloning Kit (K240020). DCL3 fragment was then introduced into pASW vector using Gateway™ LR Clonase™ II Enzyme mix (Invitrogen™ 11791021). We noticed a missing 30 bp in the DCL3 sequence compared to the AT3G43920.2 reference sequence (https://www.arabidopsis.org/servlets/TairObject?type=sequence&id=4504453036). The missing part was corrected by PCR using Oligo No. 3 and 4.

#### pASW-OsDCL3

OsDCL3a gene was cloned from the plasmid provided by National Agricultural and Food Research Organization (Clone ID J013008L07), using Oligo No. 5 and 6 and then assembled into pASW vector by NEBuilder Hifi DNA Assembly kit (NEB).

#### pASW-OsDCL5

Four DNA fragments of the OsDCL5 ORF (199–927 nt, 912–2478 nt, 2389–4785 nt and 4786–4914 nt) were amplified from *Oryza Sativa* anther cDNA using Oligos No. 7–14. A DNA fragment corresponding to 1–198 nt of OsDCL5 ORF could not be amplified from anther cDNA, and was therefore synthesized by PCR using Oligos No. 15–19 according to the reference sequence of OsDCL3b (CDS) from rap-db (The Rice Annotation Project Database) (https://rapdb.dna.affrc.go.jp/viewer/gene_detail/irgsp1?name=Os10t0485600-01;feature_id=339409). These five DNA fragments (1–198 nt, 199–927 nt, 912–2478 nt, 2389–4785 nt and 4786–4914 nt) were assembled into the pASW vector using NEBuilder Hifi DNA Assembly kit (NEB).

#### PAZ exchange plasmids

DNA fragments comprising 2560–3030 nt of OsDCL3 and OsDCL5 ORFs were exchanged between the two genes. The plasmids (pASW-OsDCL3_PAZ5 and pASW-OsDCL5_PAZ3) were prepared by PCR using Oligos No. 20–27 and the NEBuilder Hifi DNA Assembly kit (NEB).

### Cell culture

*Drosophila* Schneider 2 cells (S2 cells) were cultured in Schneider’s *Drosophila* Medium (Gibco) supplemented with 10% (v/v) Fetal Bovine Serum (FBS) (Sigma) and antibiotics at 28 °C, sealed with parafilm.

### Production of SBP-tagged AtDCL3, OsDCL3, OsDCL5 proteins in *Drosophila* S2 cells

S2 cells (1–1.5 × 10^7^ cells/10 cm dish) were transfected with 10 µg pASW plasmids carrying plant DCL3 family genes (pASW-AtDCL3, OsDCL3 or OsDCL5) with 20 µl X-tremeGENE™ HP DNA Transfection Reagent (Roche) following manufacturer’s instructions. The transfected cells were harvested after 72 hours for lysate preparation.

### Cell lysate preparation

S2 cells were harvested by centrifugation using a swinging-bucket rotor at 1,500 × *g* for 3 min at room temperature. The cell pellet was washed by cold PBS (pH 7.4) and was centrifuged at 1,500 × *g* at 4°C. The pellets were then weighed and resuspended in equal volumes of Hypotonic buffer [10 mM HEPES-KOH (pH 7.4), 10 mM potassium acetate, 1.5 mM magnesium acetate] containing 5 mM dithiothreitol (DTT) and 1 × EDTA-free Complete Protease Inhibitor tablets (Roche) by tapping and inverting the tubes. The suspension was incubated on ice for 15 minutes, and then mixed thoroughly with a vortex mixer. A cell disruption vessel (Parr Instrument Company) was used to break open the cells. The lysate was clarified by centrifugation at 17,000 × *g* for 20 min at 4°C. The supernatant was flash frozen in liquid nitrogen and immediately stored at −80°C in single-use aliquots.

### Protein purification by Streptavidin beads

Streptavidin Sepharose High Performance beads (GE Healthcare), equivalent to 25% of the lysate by volume, were washed with 1 ml lysis buffer [30 mM HEPES-KOH (pH 7.4), 100 mM potassium acetate, 2 mM magnesium acetate], and then mixed gently with the lysate. The suspension was incubated for 1 hour at 4°C on a rotator and then washed three times with wash buffer (1 × lysis buffer containing 800 mM NaCl and 1% (v/v) Triton X-100). The SBP-tagged protein was then eluted with biotin elution buffer (1 × lysis buffer, 5 mM DTT, 30% glycerol and 2.5 mM biotin) at 4°C on a rotator for 20 minutes, and the elution step repeated 3 times. The eluates were flash frozen in liquid nitrogen and immediately stored at −80°C in single-use aliquots after adding BSA to a final concentration of 0.2 mg/ml.

### Preparation of radiolabeled dsRNA substrates

The sequences of the sense and antisense RNAs used in this study are following: 38-nt sense strand, GAAUUGCUCAACAGUAUGGGCAUUUGACGCAGCCUCCC; blunt (antisense), GGGAGGCUGCGUCAAAUGCCCAUACUGUUGAGCAAUUC; 1ovr (antisense), GGAGGCUGCGUCAAAUGCCCAUACUGUUGAGCAAUUCC; 2ovr (antisense), GAGGCUGCGUCAAAUGCCCAUACUGUUGAGCAAUUCAC. Single-stranded RNAs with a 5′ hydroxyl group (OH) were synthesized by GeneDesign Inc.(Osaka Japan), while the sense strand RNA with a 5′ triphosphate was synthesized by Bio-Synthesis (Texas, USA). The antisense strand with a 3′ phosphate was radiolabeled by T4 Polynucleotide Kinase (3’ phosphatase minus) (NEB) and [γ-^32^P]ATP. Strands with a 5′ monophosphate were radiolabeled with T4 polynucleotide kinase (Takara) and [γ-^32^P]ATP. The sense and antisense strands were heat-annealed in lysis buffer as previously described^48^. The annealed dsRNAs were then separated by electrophoresis on 15% native polyacrylamide gels. The dsRNAs in gel pieces were excised and eluted by soaking in 2 × elution buffer [200 mM Tris-HCl (pH 7.5), 2 mM MgCl_2_, 300 mM NaCl, 2% SDS] overnight at room temperature. dsRNAs were mixed with glycogen and precipitated by isopropanol, then dissolved in lysis buffer.

### Dicing assay

Three nanomolar ^32^P-labeled dsRNAs and 1 or 2 nM purified recombinant proteins were incubated in 1 × lysis buffer containing 5 mM DTT, 5 mM magnesium acetate, ATP regeneration system [25 mM creatine phosphate (Sigma), 1 mM ATP, 0.03 U/µl creatin kinase (Calbiochem)], and 0.1 U/µl RNasin (Promega) at 25°C. To draw time course curves for each reaction, 2 µl of the reaction mixture was taken at 5,10, 20, 30, 45 and 60 minutes after the reaction was started, were mixed with 8 µl of low-salt PK solution [0.125% SDS, 12.5 mM EDTA, 12.5 mM HEPES-KOH (pH 7.4), 12.5% Proteinase K], and then incubated at 50°C for 10 min. An equal volume of 2 × formamide dye [10 mM EDTA (pH 8.0), 98% (w/v) deionized formamide, 0.025% (w/v) xylene cyanol, 0.025% bromophenol blue] was then added and incubated at 95°C for 2 min. The cleavage products were analyzed on 15% denaturing polyacrylamide gels and detected by autoradiography. Small RNA products were quantitated from relative band intensities measured with a Typhoon FLA 7000 image analyzer (GE Healthcare Life Sciences) and quantified using MultiGauge software (Fujifilm Life Sciences). Graphs were prepared using GraphPad Prism 8.

## Author Contributions

S.C. and H.-o.I. designed the project. S.C and W.L. performed biochemical experiments. Y.T. and H.-o.I. supervised the project. S.C. N.M, Y.T. and H.-o.I. analyzed data. S.C., Y.T. and H.-o.I. wrote the manuscript with edits provided by all authors. All authors discussed the results and approved the manuscript.

## Acknowledgements

We thank Reina Komiya for providing total RNA from the anther of *Oryza Sativa*. We also thank all the members of the Tomari laboratory for discussion and critical comments on the manuscript, and Life Science Editors for editorial assistance. S.C. gratefully acknowledges the China Scholarship Council (CSC, No. 201808050036). This work was supported by JST, PRESTO (grant JPMJPR18K2 to H.-o.I).

---

**Supplementary Figure 1:**
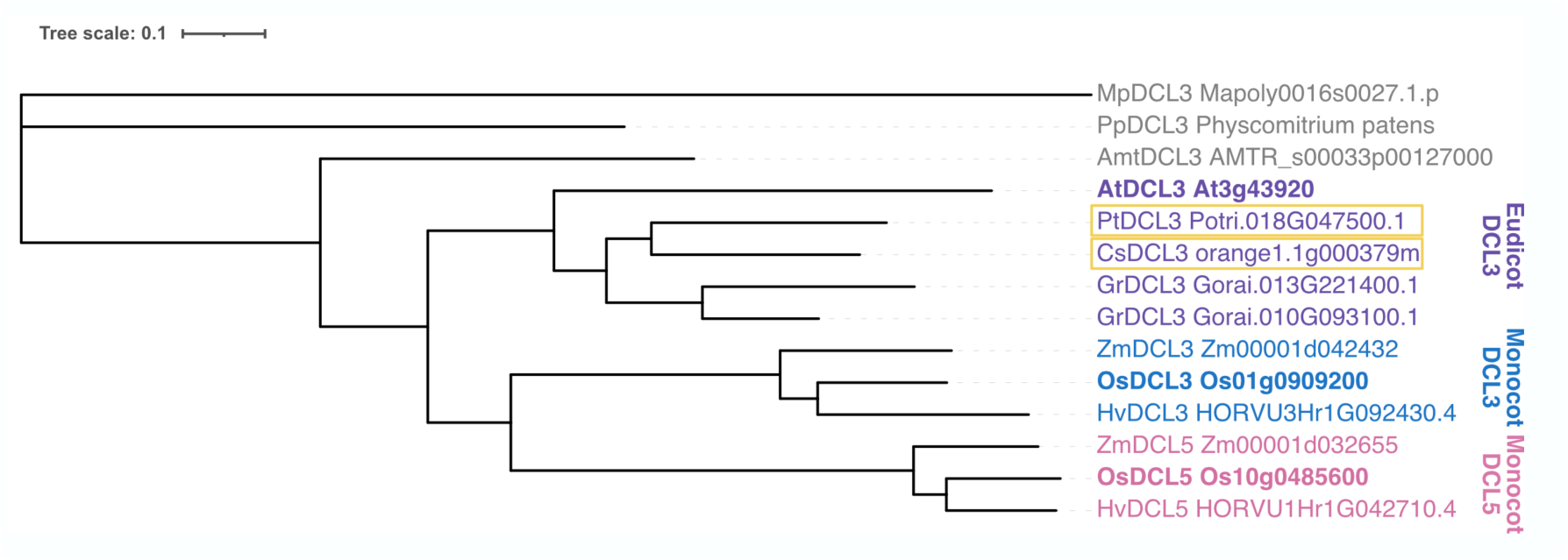
Phylogenetic tree of DCL3 family proteins. The phylogenetic tree demonstrates that eudicot DCL3, monocot DCL3 and DCL5 form separate clades. Eudicot plants that produce 24-nt reproductive phasiRNAs are marked with yellow rectangles. *Mp, Marchantia polymorpha*; *Pp*, *Physcomitrium patens*; *Amt*, *Amborella trichopoda*; *At*, *Arabidopsis thaliana*; *Pt*, *Populus trichocarpa*; *Cs*, *Citrus sinensis*; *Gr*, *Gossypium raimondii*; *Os*, *Oryza sativa*; *Zm*, *Zea mays*; *Hv*, *Hordeum vulgare*.

**Supplementary Figure 2:**
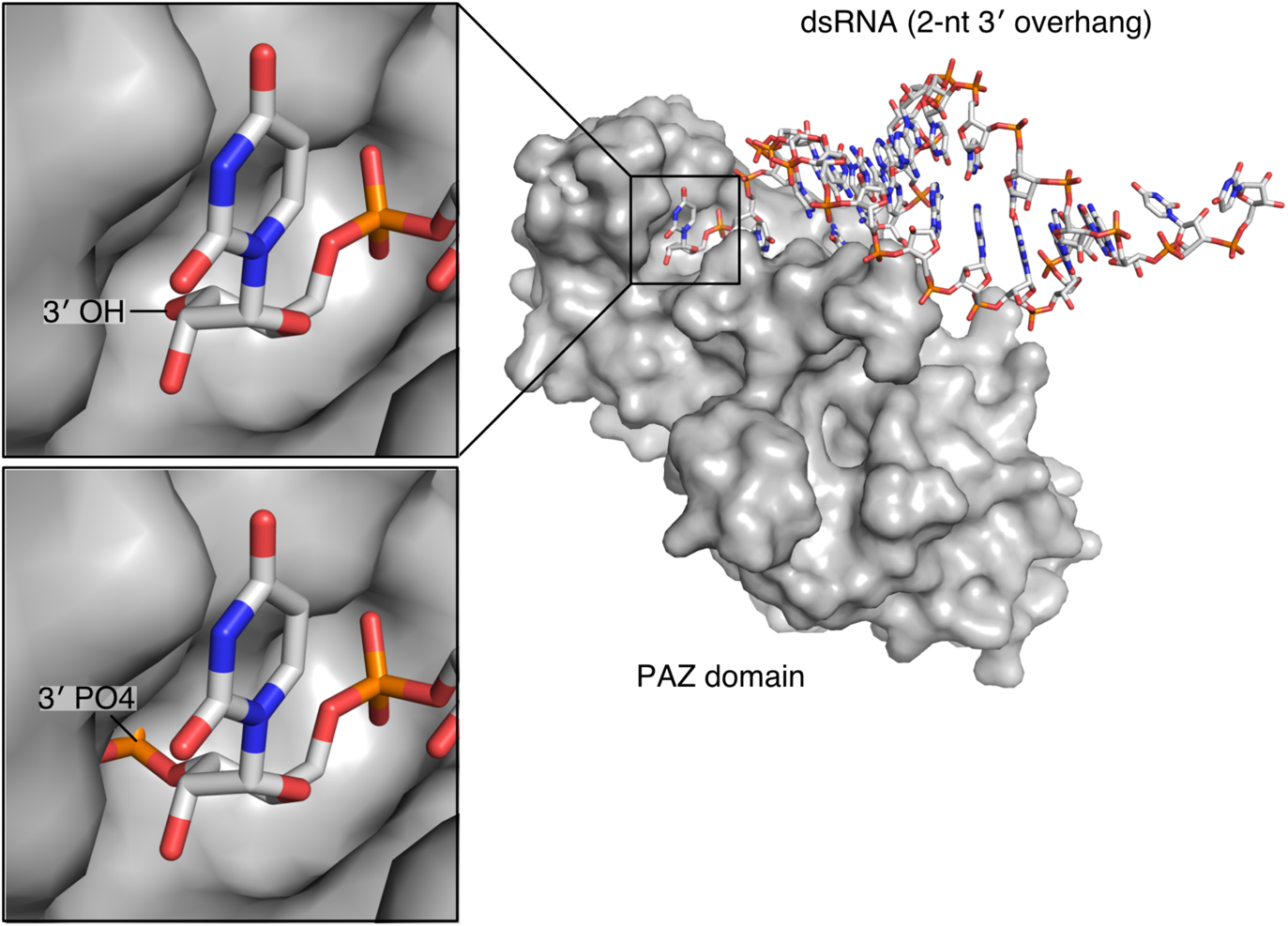
An extra 3′ phosphate blocks the binding of 1ovr substrate to the PAZ domain. Structure of the human Dicer fragment containing the PAZ domain bound to a 2-nt 3′ overhang dsRNA [Protein Data Bank (PDB) ID code: 4ngb]. The 3′ hydroxyl group of the dsRNA substrate is anchored to the 3′ binding pocket (top left). *In silico* replacement of the 3′ hydroxyl group with a 3′ phosphate group results in a steric clash with the 3′ binding pocket (bottom left).

**Supplementary Figure 3:**
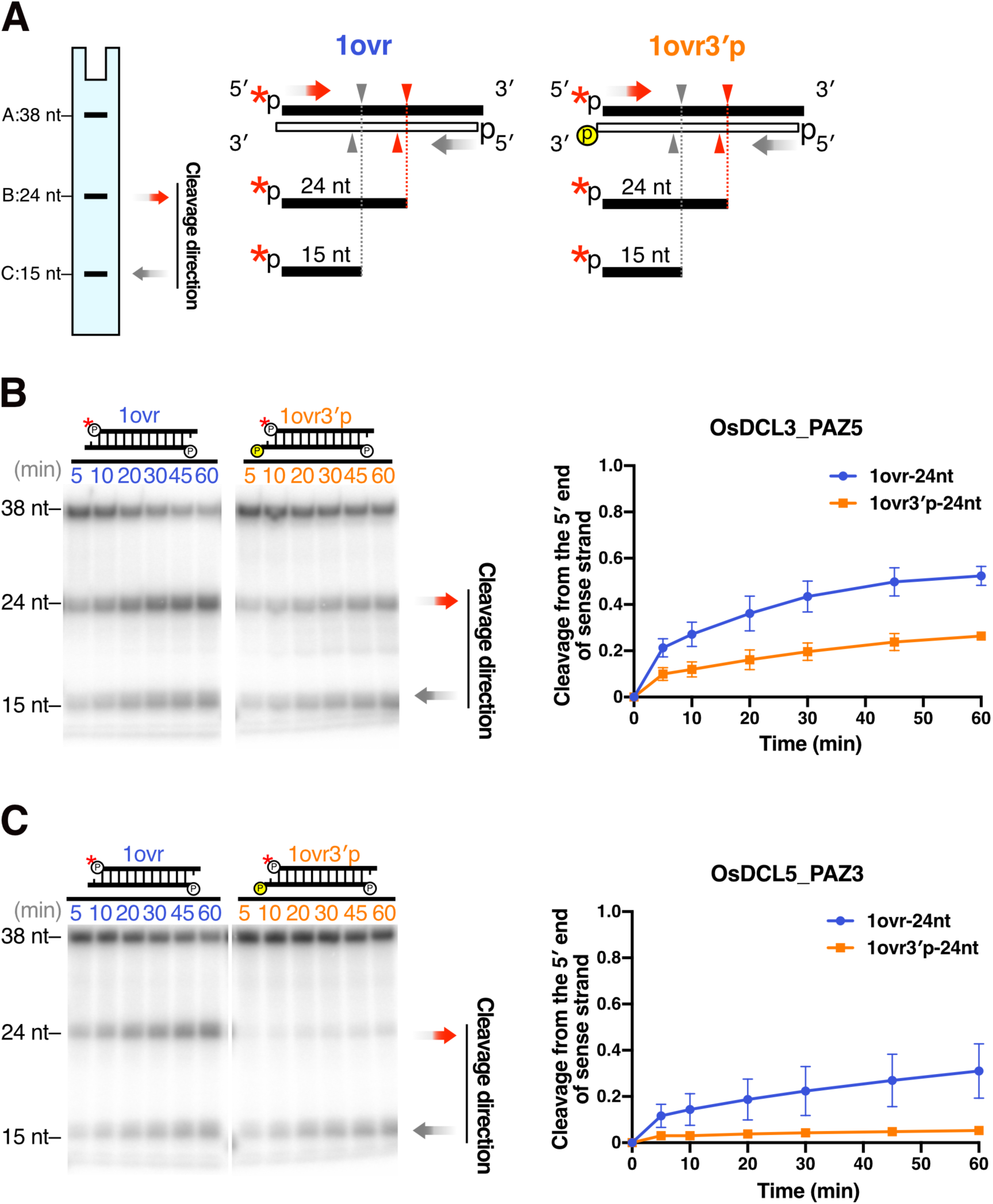
Recognition of the 3′ hydroxyl group is critical for cleavage by OsDCL5_PAZ3, but not for OsDCL3_PAZ5. (A) Dicing assays were conducted using 38-nt substrate RNAs radiolabeled on the 5′ monophosphate of sense strands with a 1ovr at both ends. An extra monophosphate group was added to the 3′ end of the antisense strand (1ovr 3′p) to test the prediction that this would disrupt recognition by the PAZ domain. Cleavage from the 5′ end of sense strands (corresponding to the 3′ phosphate on antisense strands) results in 24-nt products. (B) Left panel: A representative gel image of OsDCL3_PAZ5 dicing 1-nt overhang substrates with or without a 3′ monophosphate on the antisense strand (1ovr 3′p or 1ovr). Right panel: Quantification of dicing reaction from the 5′ end of sense strands in the left panel. Mean values ± SD from three independent experiments are shown. The extra 3′ phosphate (1ovr 3′p) moderately reduced cleavage from the 5′ end of sense strand by OsDCL3_PAZ5 compared with the 1ovr substrate. (C) Left panel: A representative gel image of dicing assay by OsDCL5_PAZ3 cleaving the 38-bp substrates 1ovr 3′p and 1ovr. Right panel: Quantification of dicing reaction from the 5′ end of sense strands in the left panel. Mean values ± SD from three independent experiments are shown. The extra 3′ phosphate (1ovr 3′p) greatly reduced cleavage from the 5′ end of sense strand by OsDCL5_PAZ3 compared with the 1ovr substrate.

**Supplementary Figure 4:**
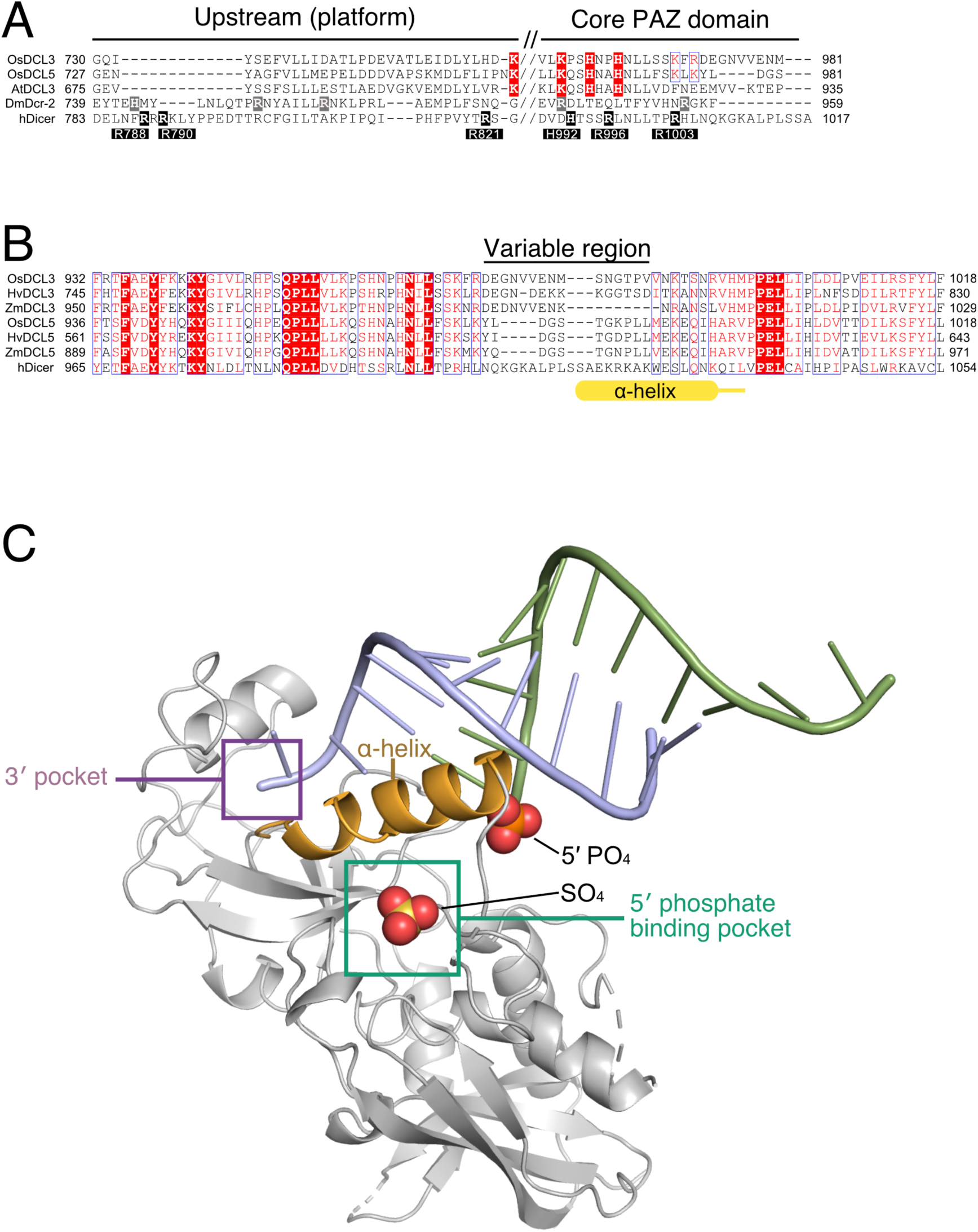
The amino-acid sequences corresponding to the *α*-helix in human DICER differ significantly between monocot DCL3 and DCL5. (A) A multiple sequence alignment of the PAZ domain of *Oryza sativa* DCL3 (OsDCL3), OsDCL5, Arabidopsis thaliana DCL3 (AtDCL3), *Drosophila melanogaster* Dcr-2 (DmDcr-2), human DICER (hDICER). White letters on black background indicate the basic amino acid residues that compose the 5′ phosphate binding pocket in human DICER. White letters on gray and red background indicate the conserved basic amino acids that correspond to the residues comprising the 5′ phosphate binding pocket of human DICER in *Drosophila* Dcr-2 and plants DCL3 family proteins, respectively. Red letters indicate basic amino acids that are not completely conserved among plant DCL3 family proteins. (B) A multiple sequence alignment of the PAZ domain of human DICER and monocot DCL3 and DCL5. *Os, Oryza sativa*; *Hv*, *Hordeum vulgare*; *Zm*, *Zea mays*. White letters on red background indicate amino acids that are perfectly conserved across human DICER and plant DCL3 family proteins. Red letters indicate amino acids that are highly conserved among human DICER and plant DCL3 family proteins. “*α-*helix” highlighted in yellow shows the position of the *α-*helix in the PAZ domain of human DICER. (C) A structure of a human DICER fragment containing the PAZ domain bound to a 2-nt 3′ overhang dsRNA (PDB: 4ngd). The structure includes the *α-*helix that separates the 5′ and 3′ binding pockets, with the 3′ end of the dsRNA anchored in the 3′ pocket and the 5′ end released from the 5′ pocket.

**Supplementary Figure 5:**
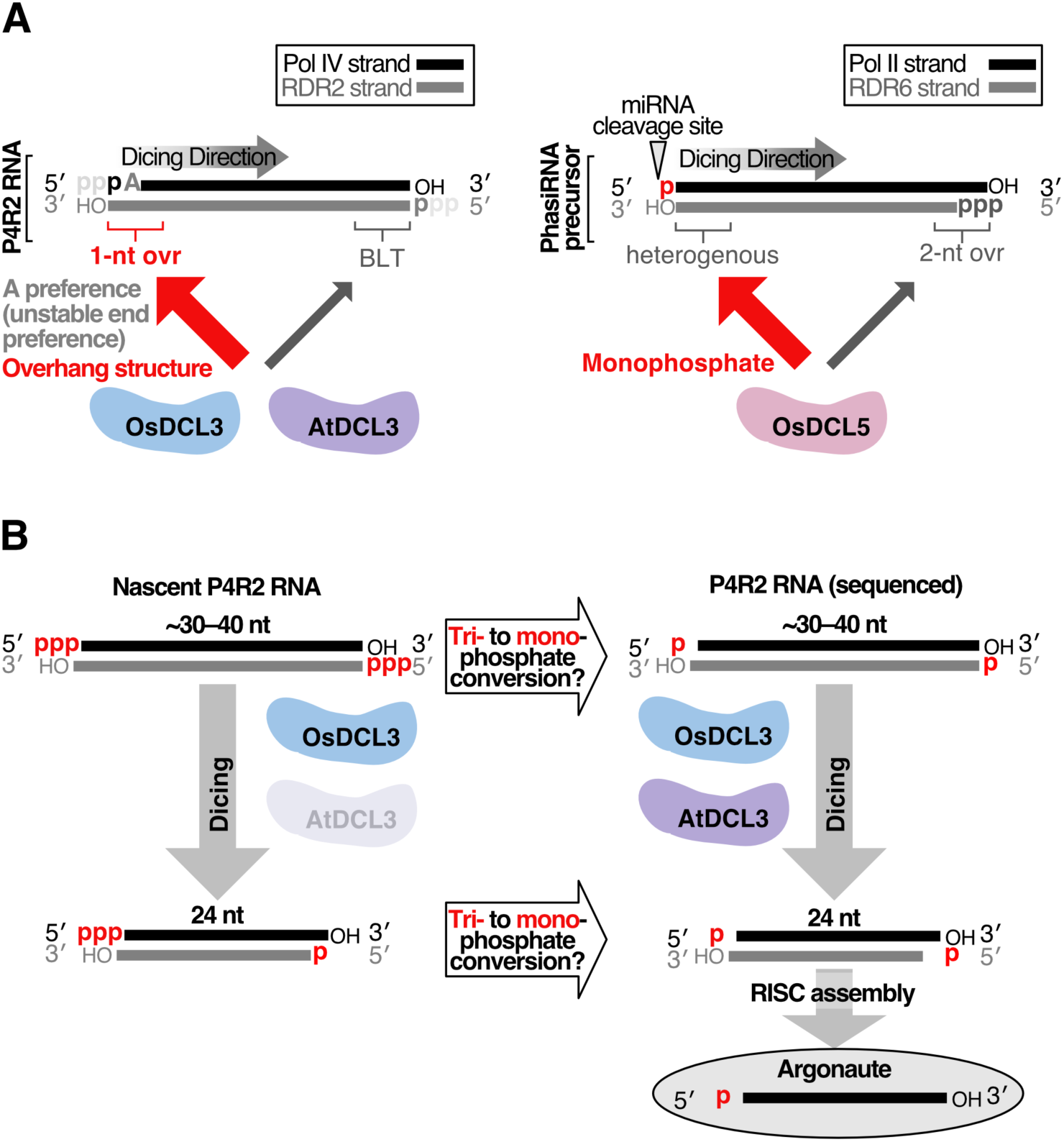
Models for unidirectional dicing by DCL3 family members and the order of tri- to monophosphate conversion and dicing by eudicot and monocot DCL3. (A) Models for unidirectional dicing mechanisms. AtDCL3 and OsDCL3 cleave P4R2 RNAs from the 5′ end of Pol IV strand as they prefer a 3′ overhang structure and unstable 5′ end. OsDCL5 cleaves the precursor from the 5′ end of the fragment due to a strong preference for the 5′ monophosphate. (B) A model for the order of tri- to monophosphate conversion and dicing by eudicot and monocot DCL3. Since AtDCL3 has a slight preference for 5′ monophosphorylated precursors, dephosphorylation prior to dicing will enhance the production of heterochromatic 24-nt siRNAs in *Arabidopsis thaliana*. In contrast, given that OsDCL3 cleaves both 5′ mono- and triphosphorylated dsRNAs with the same efficiency, OsDCL3 can effectively process P4R2 into 24-nt hc-siRNAs regardless of the tri- to monophosphate conversion.

**Table S1.**
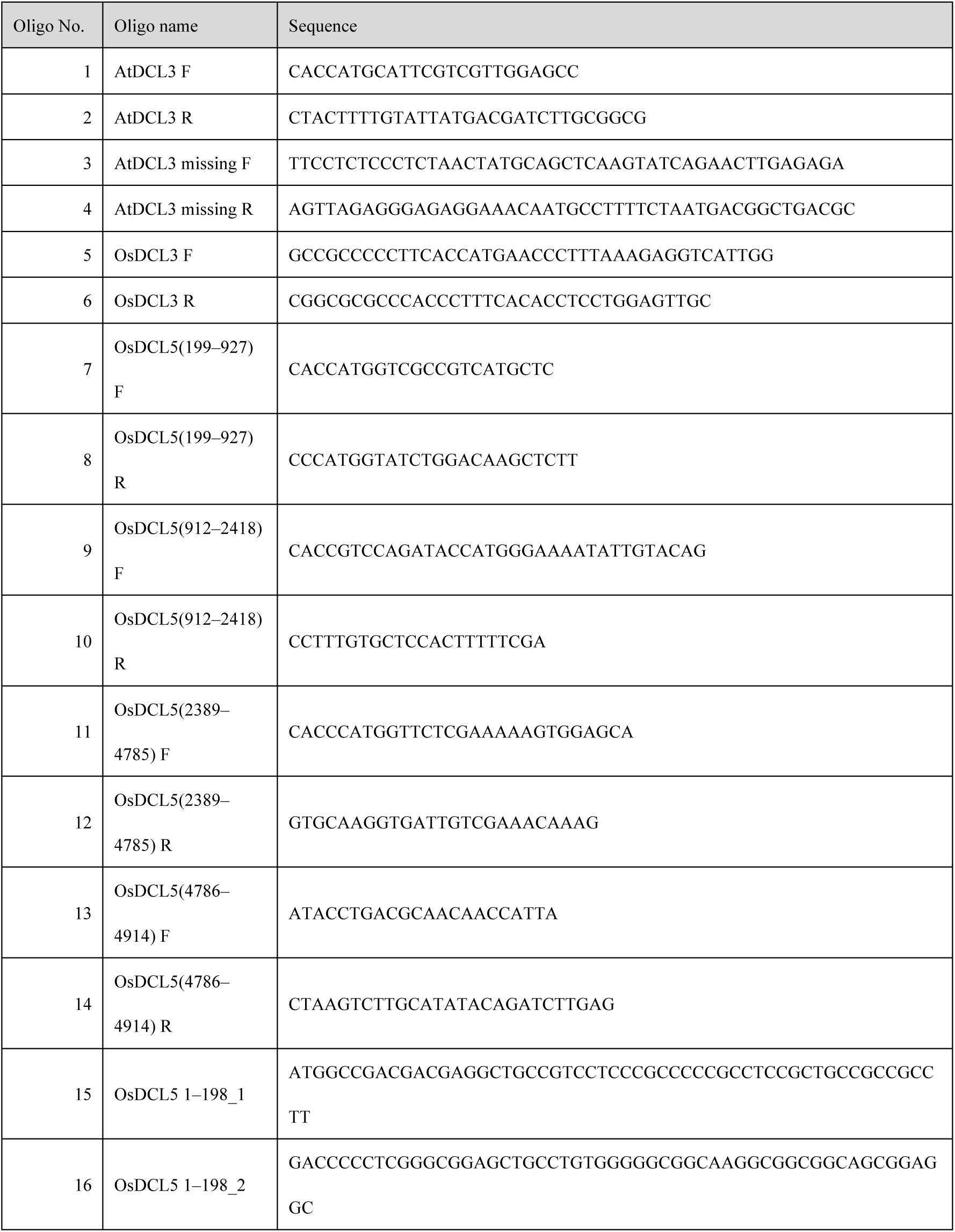
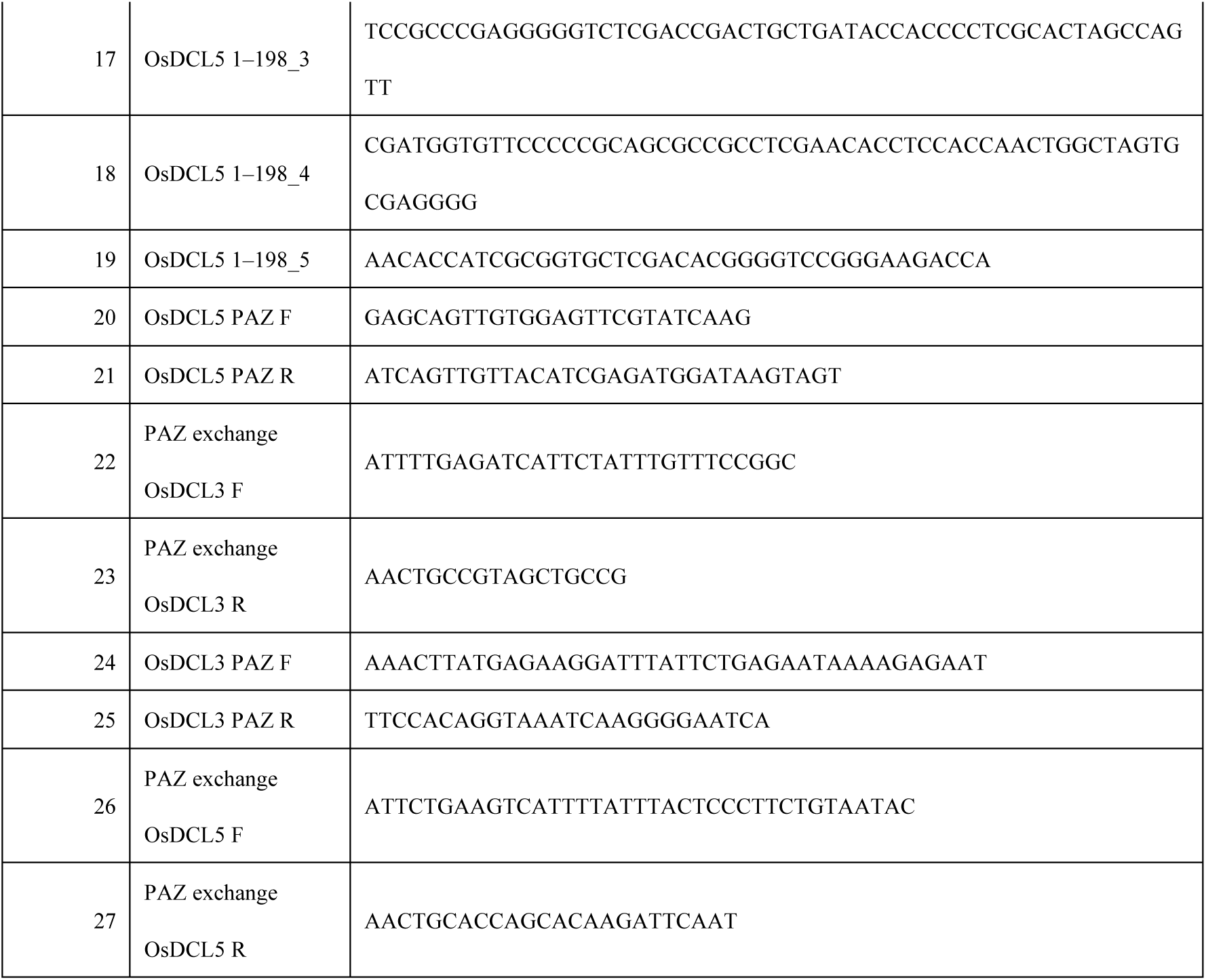
The DNA oligos used in this study.

## Supplementary Methods

### Evolutionary analysis by Maximum Likelihood method

Evolutionary history was inferred using the Maximum Likelihood method and JTT matrix-based model^1^. The tree with the highest log likelihood (−34135.57) is shown. Initial tree(s) for the heuristic search were obtained automatically by applying Neighbor-Join and BioNJ algorithms to a matrix of pairwise distances estimated using the JTT model, and then selecting the topology with superior log likelihood value. The tree is drawn to scale, with branch lengths measured in the number of substitutions per site. This analysis involved 14 amino acid sequences. There were a total of 1843 positions in the final dataset. Amino acid alignment by MUSCLE and evolutionary analyses were conducted in MEGA X^2,3^. Phylogenetic tree was modified using iTOL (https://itol.embl.de)^4^.

### Multiple alignment of the PAZ domains of animal Dicer and plant DCL3 family proteins

Multiple alignment of the PAZ domains of animal Dicer and plant DCL3 family proteins was performed using the PROMALS3D multiple sequence and structure alignment server^5^.

### Protein Structure Visualization

Protein Structure Visualization was performed by PyMol, and *in silico* replacement of the 3′ hydroxyl group on the dsRNA substrate in platform-PAZ-connector cassette (PDB ID: 4ngb) with 3′ phosphate group was done by Coot^6^.

